# Performance Attribution in pLM-Based Biological Relation Prediction

**DOI:** 10.64898/2026.06.28.735130

**Authors:** Kehan Zhu, Wenbo Zhao, Yuxuan Zhang, Zanxian Xia

## Abstract

Protein language models (pLMs) have enabled strong performance in biological relation-prediction benchmarks, yet aggregate predictive metrics do not reveal which sources of information support that performance. We examined three pLM-based case studies—MetaESI, DeepGNHV, and SAGEPhos—using frozen pLM-derived full-input baselines, component-restricted controls, train-derived endpoint or site priors, and selector-matched random restrictions.

Generic tabular learners trained on frozen pLM-derived inputs achieved strong task-specific benchmark references, reaching AUROC/AUPRC values of 0.827/0.703 for the MetaESI-derived full-input rerun, 0.922/0.708 for DeepGNHV, and 0.896/0.893 for SAGEPhos. Across tasks, substantial predictive signal remained available from restricted endpoint, site/window, motif, or train-derived prior features. In the MetaESI-derived row-level evaluation, a self-label-excluding endpoint-prior control reached AUROC/AUPRC of 0.846/0.676 without sequence embeddings, numerically close to the frozen full-input reference. In a separate frozen-pooling experiment, GARD-selected positions did not outperform count-matched random token pooling. Endpoint-cold diagnostics further showed substantial performance degradation when one or both endpoint types were unseen during training.

These findings do not diagnose leakage, imply memorization, exclude biological learning, or invalidate the evaluated models. Rather, they show that strong benchmark performance can reflect multiple overlapping information sources and that aggregate scores alone cannot establish pair-specific learning, the incremental value of task-specific architectures, selector-specific information content, or mechanistic biological interpretation. We therefore propose that such claims require matched attribution controls, representation-matched baselines, and evaluation protocols aligned with the intended generalization setting.

## 1 Introduction

Protein language models have become common upstream encoders for biological prediction, including protein–protein interaction, enzyme–substrate relation, and kinase–substrate–site prediction [5, 2]. Their pretrained representations provide a practical route to transfer information learned from large protein-sequence corpora into task-specific models. High AUROC or AUPRC on a benchmark can establish that a model is useful under a stated evaluation protocol. The same result is sometimes also used to motivate stronger interpretations about the contribution of a proposed architecture, the specificity of a predicted biological relation, or the meaning of a selected region or feature [1, 6].

Benchmark utility and performance attribution are different questions. Utility asks whether a predictor performs well on a particular test distribution. Attribution asks which information sources make that performance possible. In a biological relation benchmark, the same aggregate score may be supported by joint compatibility between two entities, properties of one endpoint, local site or motif context, repeated-entity frequency, database coverage, the construction of negative examples, classifier capacity, or a task-specific architectural operation. Standard test metrics alone do not distinguish among these possibilities.

This attribution problem is especially important when pLM embeddings are used as upstream representations. Unlike sparse handcrafted features, pLM embeddings can encode broad sequence, family, structural, and functional information learned before the downstream task is trained. A generic learner may therefore achieve strong benchmark performance from information already accessible in frozen embeddings, without implementing the proposed task-specific architecture. This does not make the proposed architecture irrelevant. Rather, without a representation-matched baseline, it is difficult to determine how much of the observed performance is attributable to the task-specific architecture rather than to information already accessible in the pretrained representation.

We use **baseline decomposition** to denote claim-matched controls that alter the information available to a predictor. A representation-matched full-input learner establishes the performance accessible without the proposed task-specific architecture. Component-restricted and train-derived prior controls test whether narrower parts of the prediction unit or repeated-entity structure already support discrimination. Matched random restrictions and increasingly stringent split or negative-sampling designs then test selector-specific and generalization claims. These controls are overlapping diagnostics rather than an additive partition.

We applied this framework to three pLM-based biological prediction settings with distinct prediction units: E3–substrate relation prediction (MetaESI[4]), host–virus protein-pair prediction (DeepGNHV[3]), and kinase–substrate–site/window prediction (SAGEPhos[7]). These case studies differ in label construction, class prevalence, representation, learner, and evaluation split, and are intended to illustrate a general attribution problem rather than to provide a cross-task ranking or a systematic survey of the field.

Our analysis asks how much benchmark performance is accessible from frozen pLM representations, individual components of the prediction unit, train-derived endpoint priors, and restriction-matched selector controls. Across three case studies, these controls reveal that strong benchmark discrimination can be supported by information sources that are not specific to the proposed task architecture.

## 2 Results

We evaluated three pLM-based biological prediction case studies that differ in prediction unit, class prevalence, representation, and evaluation protocol (Table 1). For each case study, we first established a frozen pLM-derived benchmark reference using generic tabular learners, and then compared it with controls that restrict the available information to individual components, train-derived priors, or selector-matched random restrictions.

**Table 1:** Key performance-attribution results across three pLM-based biological prediction case studies. Metrics are reported as AUROC/AUPRC on the stated held-out evaluation rows and are intended for within-task interpretation only. MetaESI frozen full-input, endpoint-prior, and GARD/random analyses are separate evidence blocks with different feature constructions and down-stream learners; they should not be interpreted as a single nested ablation chain. The DeepGNHV cold diagnostics use matched warm and endpoint-cold evaluation settings. SAGEPhos train-derived priors include exact site-window statistics. Complete source-backed metrics, additional control conditions, provenance metadata, and archived/superseded rows are provided in Supplementary Data 1.

| Case study | Evaluation setting | Frozen full-input reference (AU-ROC/AUPRC) | Key restricted or prior control (AU-ROC/AUPRC) | Selector or cold diagnostic (AUROC/AUPRC) | Attribution-relevant observation |
| --- | --- | --- | --- | --- | --- |
| MetaESI | Row-level split | 0.827 / 0.703 | Self-label-excluding endpoint prior: 0.846 / 0.676 | GARD-selected pooling: 0.819 / 0.683; random pooling: 0.820–0.823 / 0.682–0.684 | Train-derived endpoint-prior features and count-matched random token pooling supported performance numerically similar to the frozen-reference block under their respective evaluated settings. |
| DeepGNHV | Official VF1 split | 0.922 / 0.708 | Virus-only input: 0.701 / 0.213 | Cold-human: 0.830 / 0.401; cold-virus: 0.704 / 0.398; cold-both: 0.671 / 0.304 | The combined two-endpoint input was substantially stronger than either endpoint alone, while endpoint novelty reduced performance. |
| SAGEPhos | Provided test split | 0.896 / 0.893 | Site/window-only: 0.634 / 0.649; train-derived kinase/substrate/site-window prior: 0.704 / 0.700 | No source-backed cold diagnostic available | Local site/window information and train-derived site-inclusive prior features retained substantial predictive signal. |

### 2.1 Frozen pLM-derived full-input baselines establish strong benchmark references

We first quantified the performance accessible to frozen pLM-derived inputs without training the corresponding task-specific neural architectures (Fig. 1). Under the MetaESI row-level split, a verified full-input ESM2 mean-embedding rerun coupled to AutoGluon reached AUROC 0.827 and AUPRC 0.703. The test-set positive prevalence was 0.333. For DeepGNHV, a ProtT5 full-input baseline using both endpoint embeddings and pairwise feature operators reached AUROC/AUPRC 0.913/0.690 on the official VF1 split. A simpler two-endpoint concatenation of the human and viral embeddings performed slightly better, reaching 0.922/0.708; the test prevalence was 0.091. For SAGEPhos, a combined site-conditioned ESM2 feature set comprising kinase, substrate, and phosphorylation-site/window information reached 0.896/0.893 on the provided test split, whose positive prevalence was 0.503.

**Figure 1:**
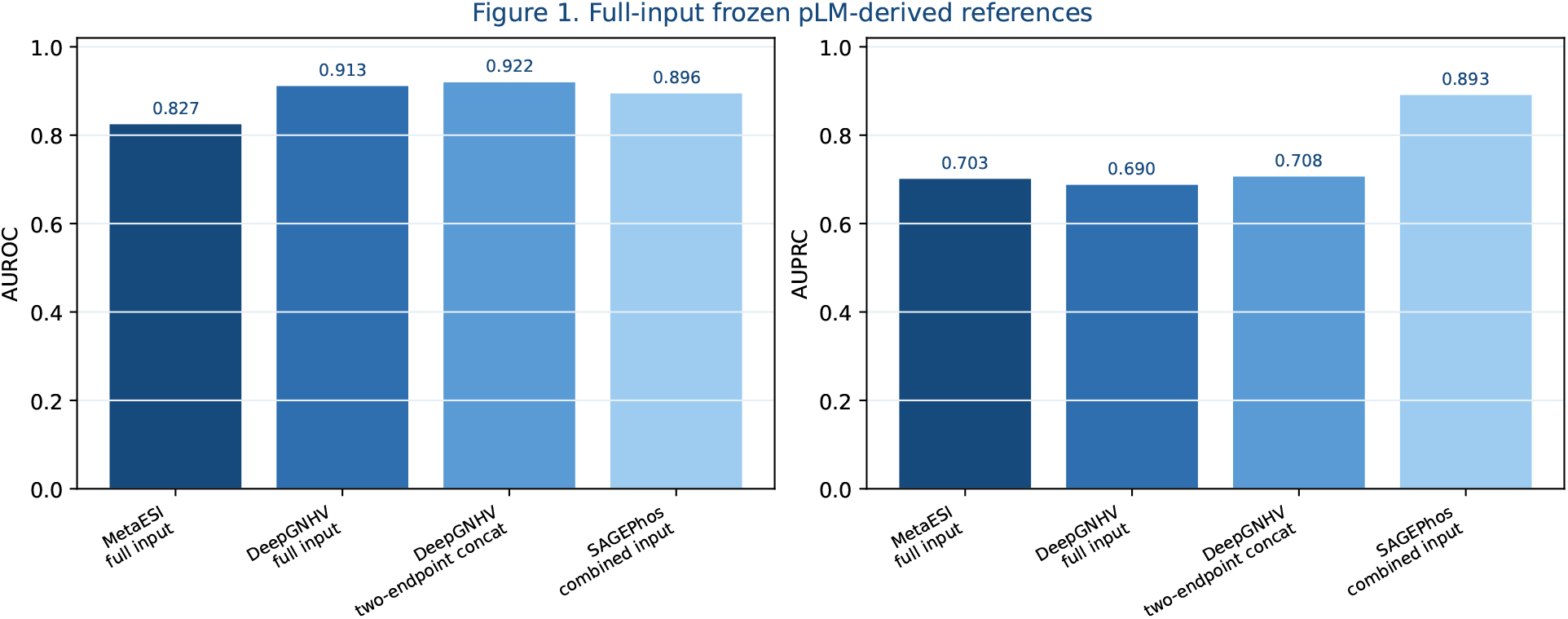
Full-input frozen pLM-derived benchmark references across three biological prediction case studies. Generic tabular learners trained on frozen pLM-derived full-input features achieved strong task-specific reference performance for the MetaESI-derived E3–substrate evaluation, DeepGNHV human–virus protein-pair evaluation, and SAGEPhos kinase–substrate– site/window evaluation. Left, AUROC; right, AUPRC. Metric values are shown above bars. These values are presented as within-task benchmark references only and are not intended for direct comparison across case studies, which differ in prediction unit, class prevalence, representation, split definition, and downstream training procedure.

These results are not intended as cross-task comparisons: the three benchmarks differ in prediction unit, prevalence, split, representation, and downstream training procedure. They establish task-specific reference points. In each case, strong benchmark performance was accessible from frozen pLM-derived inputs and generic tabular learning before any performance was assigned to the original task-specific architecture.

### 2.2 Restricted components and train-derived priors retain substantial predictive signal

We next evaluated component-restricted inputs and train-derived prior controls while retaining the corresponding benchmark rows and split definitions (Fig. 2). In the MetaESI endpoint-ablation block, concatenating the E3 and substrate embeddings without the additional absolute-difference and element-wise-product operators reached AUROC/AUPRC 0.831/0.688. The endpoint-only controls were asymmetric: E3-only reached 0.669/0.404, whereas substrate-only reached 0.478/0.301. Because these rows were generated in an earlier ablation block, they are interpreted relative to one another rather than as a strict nested comparison with the full-input reference reported here.

**Figure 2:**
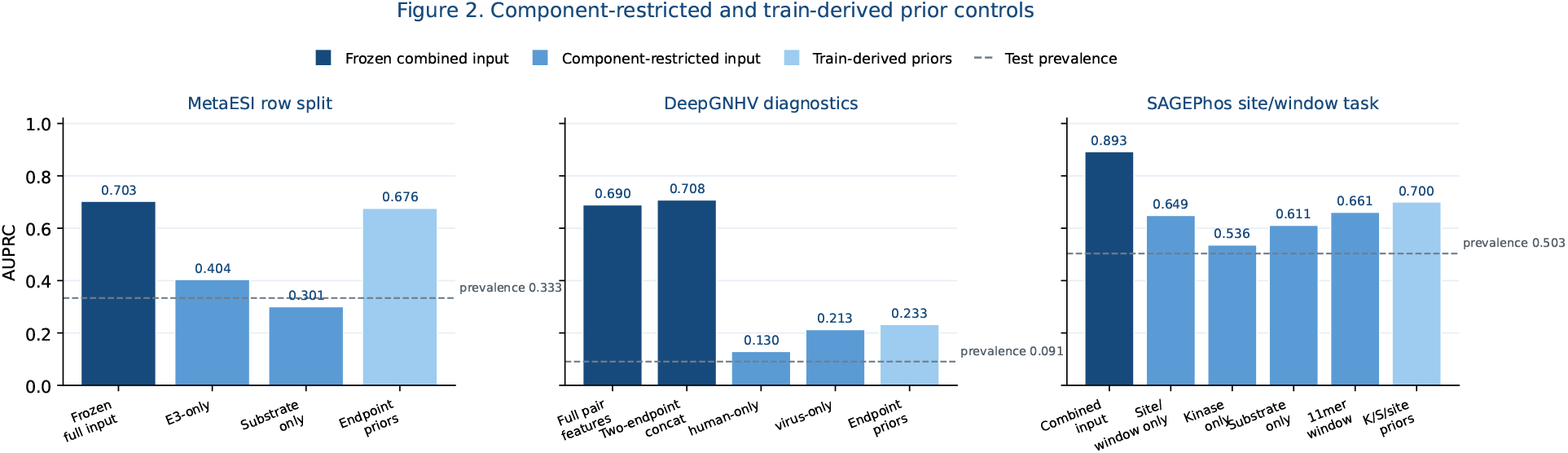
Component-restricted and train-derived prior controls retain task-dependent predictive signal. AUPRC is shown for frozen combined-input references, component-restricted inputs, and clean train-derived prior controls across the three case studies. Dashed horizontal lines indicate the positive prevalence in the evaluated test set. Left, MetaESI-derived row-level evaluation: E3-only and substrate-only frozen ESM2 controls are shown alongside a self-label-excluding endpoint-prior control. Middle, DeepGNHV diagnostics: human-only and virus-only ProtT5 controls are shown together with a clean endpoint-prior diagnostic evaluated on the original VF1 catalog. Right, SAGEPhos site/window task: site/window-only, kinase-only, substrate-only, local 11-mer window, and kinase/substrate/site-window prior controls are shown. Because feature sources and, for some supplementary diagnostics, evaluated row populations differ, the figure is intended for performance attribution within each task rather than direct estimation of numerical architecture contributions.

DeepGNHV showed a different endpoint pattern. Human-only and virus-only ProtT5 controls reached AUROC/AUPRC 0.550/0.130 and 0.701/0.213, respectively, compared with 0.922/0.708 for the combined-input reference. The viral endpoint therefore retained more benchmark-accessible signal than the human endpoint under the official VF1 split, although neither endpoint-only control approached the full two-endpoint input.

For SAGEPhos, the sample unit includes the kinase, substrate, and phosphorylation site or local sequence window. Site/window-only, kinase-only, and full-substrate-only controls reached AUROC/AUPRC 0.634/0.649, 0.547/0.536, and 0.631/0.611, respectively, compared with 0.896/0.893 for the combined reference. Non-pLM local-sequence controls were also predictive: an 11-residue one-hot window reached AUPRC 0.661, amino-acid property features reached 0.645, and local k-mer composition reached 0.640. Because local sequence context is biologically integral to kinase specificity, these controls are interpreted as component-level attribution diagnostics rather than as evidence against the relevance of site or motif information.

### 2.3 Self-label-excluding endpoint priors approach the MetaESI row-split reference

Train-derived degree and frequency controls were analyzed separately from the pLM feature ablations because they use different feature spaces and, in MetaESI, different downstream learners. For the clean MetaESI control, the features comprised endpoint occurrence counts, positive counts, Laplace-smoothed empirical positive rates, unseen-endpoint indicators, and simple cross-endpoint interactions. Training-row features were generated by inner out-of-fold estimation with explicit self-label exclusion; validation and test features were derived from the outer training set only.

The primary inner-out-of-fold LogisticRegression control reached AUROC 0.846 and AUPRC 0.676. Relative to the frozen full-input reference, its AUROC was numerically 0.019 higher and its AUPRC was 0.027 lower. A leave-one-edge-out LogisticRegression sensitivity reached 0.849/0.692, while an inner-out-of-fold fixed HistGradientBoosting sensitivity reached 0.847/0.684 (Table S1). None of these controls used protein sequence embeddings or explicit pairwise sequence-comparison features. The values should not be interpreted as statistical equivalence to the full reference, but they show that train-derived endpoint-prior features can yield a numerically similar result under the row-level split.

Clean supplementary prior diagnostics were also evaluated for DeepGNHV and SAGEPhos, but these were not designed as directly matched architecture-contribution estimates. For DeepGNHV, a clean full-catalog human/virus endpoint-prior LogisticRegression diagnostic reached AUROC/AUPRC 0.706/0.233 on the original VF1 catalog. This diagnostic is not directly comparable with the frozen pLM endpoint block because the evaluated row populations differ. For SAGEPhos, clean kinase/substrate/site-window priors reached 0.704/0.700. These priors include exact 11-mer/site-window statistics, so local site frequency is itself part of the feature source.

These controls do not determine why endpoint priors are predictive. Biological hubness or substrate propensity, experimental attention, database coverage, repeated sampling, negative-set construction, and label-construction processes may all contribute, and these explanations are not mutually exclusive. These controls therefore identify a source of benchmark-accessible information without determining its biological or data-generating origin.

### 2.4 GARD-selected pooling does not outperform count-matched random pooling

We next considered a narrower, model-specific question in MetaESI: whether the identity of GARD-selected residues contributed more predictive information than an equally sized restriction of the frozen ESM representation (Fig. 3). The comparison was performed in a fixed frozen-pooling LightGBM pipeline and is separate from the AutoGluon full-input rerun, manual endpoint-ablation ensemble, and clean endpoint-prior blocks.

**Figure 3:**
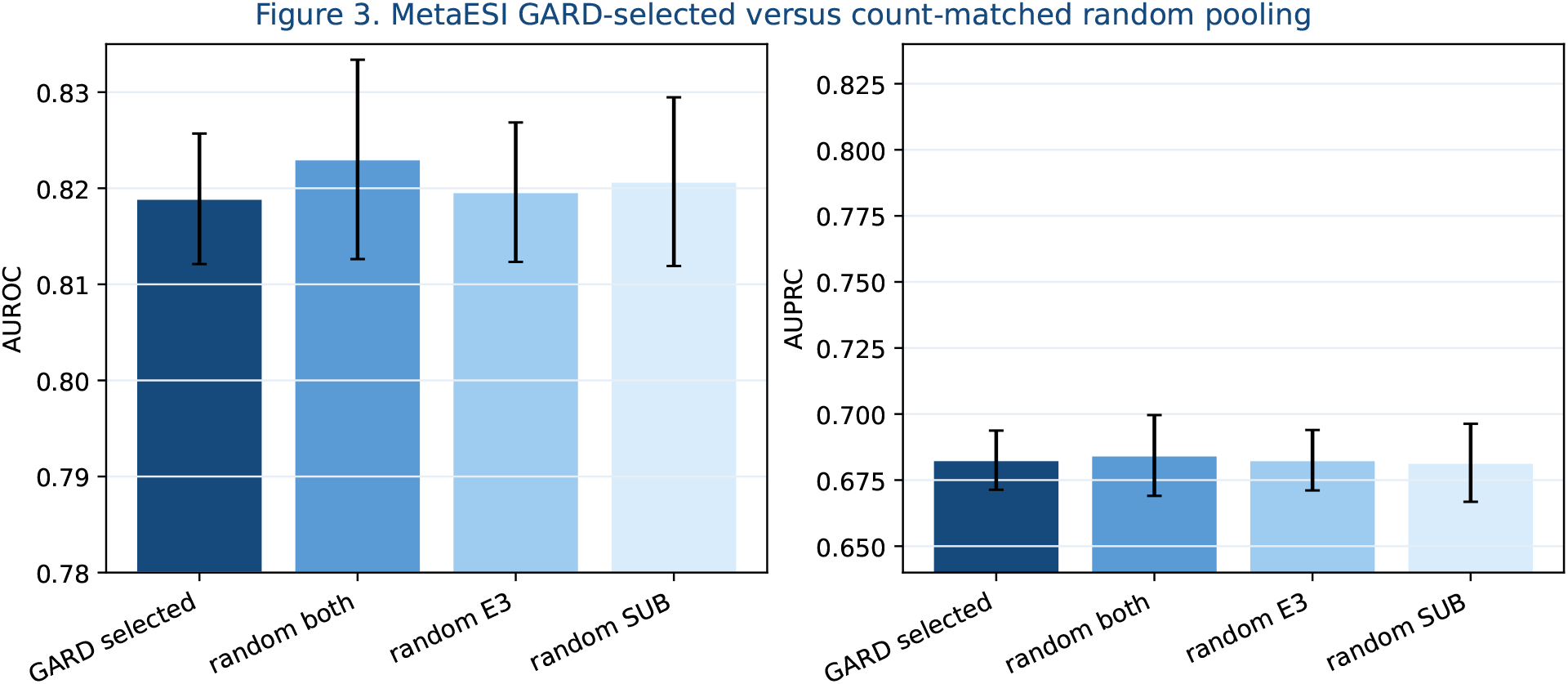
GARD-selected pooling does not outperform count-matched random token pooling in the evaluated frozen-token LightGBM pipeline. Frozen ESM2 token representations were pooled over GARD-selected positions or over random positions matched for retained token count and sampled from both endpoints, the E3 endpoint, or the substrate endpoint. Error bars show the standard deviation reported in the source summary: five fold-level evaluations for GARD-selected pooling and 10 random seeds *×* five folds for each row-level random-control condition. The y-axes are truncated for visual resolution; conclusions are based on point estimates and source-summary variability, not visual bar height. Under the tested row-level evaluation, the observed point estimates did not show an advantage for GARD-selected pooling over count-matched random pooling. This comparison tests the information content of selected token identities within a fixed pooling and LightGBM readout, rather than the complete MetaESI architecture or interactions between GARD and downstream neural modules.

Under row-level evaluation, GARD-selected pooling reached AUROC/AUPRC of 0.819/0.683. Count-matched random pooling from both endpoints reached 0.823/0.684, while random pooling from the E3 and substrate sides reached 0.820/0.683 and 0.821/0.682, respectively. Across row-level, E3-cold, and substrate-cold evaluations, the observed point estimates did not show a consistent advantage for GARD-selected positions over count-matched random pooling (Fig. S3C; Table S3).

The selector retained broad and, on the substrate side, frequently discontinuous sequence coverage rather than uniformly isolated residue-scale regions (Fig. S3). Because ESM2 token embeddings are contextual representations [5], retaining a token should not be interpreted as isolating information attributable only to that residue or its immediate local state. This comparison does not isolate the complete MetaESI architecture or interactions between GARD and downstream neural modules; it tests whether selected token identity adds predictive value beyond a matched restriction of the frozen representation.

### 2.5 Endpoint-cold diagnostics delimit the current evidence

Endpoint-cold diagnostics were available for MetaESI and DeepGNHV (Fig. S1; Table S2). For MetaESI, an earlier feature-ablation diagnostic artifact reported AUROC/AUPRC of 0.798/0.651 under E3-cold evaluation and 0.820/0.684 under substrate-cold evaluation for the full frozen-representation baseline. Because these runs preceded the full-input reference reported here, they are presented as supplementary endpoint-novelty diagnostics rather than as directly matched estimates of performance degradation relative to the current warm evaluation. The endpoint-restricted controls also changed in a direction-dependent manner, indicating that the apparent difficulty of a cold split depends on which endpoint is withheld and which endpoint-level information remains available to the predictor.

For DeepGNHV, the two-endpoint concatenation baseline was evaluated under matched warm and endpoint-cold conditions. Its warm AUROC/AUPRC of 0.922/0.708 decreased to 0.830/0.401 for cold-human, 0.704/0.398 for cold-virus, and 0.671/0.304 for cold-both evaluation. Test-set positive prevalences were 0.088, 0.115, and 0.111, respectively. Performance remained above prevalence-based baselines in all cold settings, but decreased substantially, with the largest reduction when both endpoint types were unseen during training.

These analyses are endpoint-novelty stress tests rather than hard-negative, matched-negative, family-aware, homology-aware, or external-cohort evaluations. No source-backed hard- or matched-negative evaluation was available across the three case studies, and no kinase-, substrate-, or site-cold diagnostic was established for SAGEPhos. Accordingly, the present study supports performance-attribution diagnostics under the evaluated benchmark settings: commonly reported benchmark performance can be substantially shaped by representation-accessible and endpoint-level signal. It does not establish robustness to family-level novelty, hard negatives, or external biological generalization.

### 2.6 Multiple signal sources can coexist with strong benchmark performance

Taken together, the case studies show that strong benchmark performance can be supported by multiple, overlapping sources of predictive information. The clearest example was the MetaESI-derived downstream evaluation setting used here, in which an endpoint-prior control constructed without using the held-out row label achieved an AUROC slightly above, and an AUPRC close to, the frozen full-input reference under row-level evaluation. This result does not imply leakage, memorization, or the absence of biologically meaningful signal. Rather, it shows that this evaluated row-level benchmark contains substantial endpoint-prior structure, such that its aggregate score alone cannot identify how much performance reflects pair-specific recognition, incremental value from the task-specific architecture, or endpoint-associated information already available from the training data.

DeepGNHV and SAGEPhos likewise exhibited substantial endpoint-, site/window-, motif-, and train-derived prior signal, although the magnitude, evaluated row population, and biological interpretation differed across tasks. Strong performance should therefore be interpreted as an aggregate outcome rather than as direct evidence for any single proposed mechanism. Performance attribution requires controls matched to the claim being made: endpoint-prior controls for pair-specificity claims, representation-matched baselines for architecture-necessity claims, and independent validation for mechanistic or biological-interpretability claims.

## 3 Discussion

The central result of this study is not that strong benchmark performance is unreal, but that it is underdetermined. In pLM-based biological relation prediction, the same AUROC or AUPRC can be supported by information already encoded in a pretrained representation, endpoint propensity, local sequence or motif context, repeated-entity and database-frequency structure, selector-induced restriction, and task-specific modeling of the joint relation. Aggregate benchmark metrics establish predictive utility under a specified data distribution; they do not decompose why that utility is achieved.

The MetaESI-derived row-level evaluation provides the clearest illustration. A self-label-excluding endpoint-prior LogisticRegression control reached AUROC 0.846 and AUPRC 0.676 without sequence representations, numerically close to the frozen ESM2 full-input reference. This is not a matched estimate of the original architecture’s contribution, because the two analyses used different learners and training procedures. It nevertheless shows that the evaluated row split strongly rewards train-derived endpoint-frequency structure. Under such conditions, a strong score cannot by itself distinguish pair-specific sequence compatibility, architecture-specific integration, or endpoint-associated information already available from the training data.

This does not make endpoint, motif, site-window, or frequency signal illegitimate. These features may reflect genuine biology, including enzyme specificity, substrate context, family structure, and local biochemical constraints. They may also reflect ascertainment, experimental attention, database coverage, repeated sampling, negative-set construction, and other properties of the data-generating process. The relevant question is therefore not whether a simple control is “biological” or an “artifact,” but whether the reported performance requires the full relation and the proposed architectural operation beyond information already accessible from a restricted component of the prediction unit.

This distinction is particularly important when frozen pLM representations are used upstream. Generic tabular learners, including high-capacity systems such as AutoGluon, are not mechanistically interpretable models. Their role here is diagnostic: they estimate how much predictive signal can be extracted from the same frozen representation without task-specific biological architecture. If a specialized downstream architecture substantially exceeds such a reference under matched evaluation, its additional inductive bias may be empirically justified. If it does not, comparable predictive performance alone cannot establish the necessity of the specialized architecture or the biological meaning of its internal operations. This matters practically as well as epistemically: architecture-specific pipelines impose additional implementation complexity, optimization burden, computational cost, and interpretive burden. These costs are most defensible when accompanied by demonstrable predictive or independently validated biological gains.

The selector analysis extends this principle to individual architectural components. Across row-level, E3-cold, and substrate-cold evaluations within a fixed frozen-pooling LightGBM pipeline, GARD-selected positions did not show a reproducible predictive advantage over count-matched random positions sampled from the E3, substrate, or both endpoints. This does not invalidate the biological relevance of GARD-defined regions or the complete MetaESI architecture. Rather, it shows that selector-specific claims require a restriction-matched reference. When a model isolates residues, tokens, local windows, latent dimensions, or subgraphs, the corresponding random control should preserve the relevant information budget, including token count, sequence coverage, dimensionality, graph size, and endpoint allocation.

This requirement is especially important for pLM-derived token representations. Because ESM2 token embeddings are contextualized by the full sequence, a retained token need not represent only information localized to that residue. Moreover, the GARD-retained substrate representation commonly comprised multiple fragments, including broad fragments spanning tens to hundreds of residues. The operation should therefore be interpreted primarily as contextual representation filtering rather than direct residue-level localization unless supported by independent residue-level validation.

The original studies included architecture-level analyses such as module ablations, perturbation tests, cold-start evaluations, and selected biological case studies. Our analysis addresses a different question: whether benchmark-level performance can be attributed to the proposed architecture after accounting for frozen-representation signal, restricted components, train-derived priors, and selector-matched controls. Architecture-level ablation tests whether a component improves a specified pipeline; benchmark-level attribution tests whether the observed score identifies the source of that improvement. Both are needed when predictive performance is used to support biological interpretation.

These observations motivate claim-matched evaluation. Architecture-contribution claims require representation-matched reference learners evaluated on the same rows and upstream features. Relation-specific claims require endpoint-, site-, local-context-, and train-derived-prior controls, whereas selector-specific claims require matched random restrictions. Claims about novelty or biological generalization require progressively stricter evaluation, including cold splits, family- or homology-aware partitions, matched negatives, external validation, and, where mechanism is claimed, orthogonal biological evidence. Such controls are protective rather than adversarial: strong performance after them would strengthen the corresponding interpretation.

Several limitations bound the present study. The three case studies are heterogeneous and are not intended as a cross-task ranking or systematic survey. AutoGluon is an architecture-agnostic but potentially high-capacity reference; we did not reimplement every original model or equalize optimization budgets. The MetaESI frozen-pLM, endpoint-prior, and GARD/random analyses are separate evidence blocks rather than a nested ablation chain. DeepGNHV prior diagnostics use a different evaluable row population from the frozen-pLM endpoint controls, and SAGEPhos priors include exact site-window statistics. Cold evaluation remains incomplete, and hard-negative, family-aware, homology-aware, and external evaluations were unavailable. The study therefore distinguishes benchmark performance from performance attribution; it does not quantify the contribution of every original architecture or determine the biological origin of all predictive signal.

## 4 Methods

### 4.1 Study design and scope

We conducted a source-audited baseline-decomposition study using verified artifacts from three pLM-based biological relation-prediction projects. The analysis was retrospective: we did not attempt a complete reimplementation of each original architecture. Instead, we assembled full-input frozen-representation baselines and progressively restricted controls designed to test distinct attribution questions. Evidence blocks that used different representations, downstream learners, or training procedures were kept separate and were not treated as a single nested ablation. The synchronized result table, coverage matrix, figure source tables, configurations, feature manifests, split manifests, predictions, and provenance index were generated from the audited repository artifacts.

### 4.2 Case studies and prediction units

MetaESI[4] was treated as E3–substrate binary relation prediction under a row-level split. The verified split contained 5,533 training rows, 791 validation rows, and 1,581 test rows, with positive prevalences of 0.333, 0.334, and 0.333, respectively. DeepGNHV[3] was treated as host–virus protein-pair prediction under the official VF1 split; the test set contained 40,533 pairs with positive prevalence 0.091. SAGEPhos[7] was treated as kinase–substrate–site/window prediction rather than a pure protein-pair task. Its verified split contained 29,375 training rows, 3,672 validation rows, and 3,672 test rows, with test prevalence 0.503. Exact split manifests, training-set counts for each pipeline, and endpoint-cold split summaries are provided in the supplementary source tables.

### 4.3 Frozen pLM representations and full-input baselines

For MetaESI, whole-protein ESM2-derived mean embeddings were used for the E3 and substrate. The full-input representation was

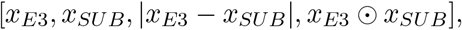

where ⊙ denotes the element-wise product. The current MetaESI reference was obtained from a verified row-level rerun using the full frozen representation. Two-endpoint, E3-only, and substrate-only controls were obtained from a separate endpoint-ablation block. The two-endpoint control retained the concatenated E3 and substrate embeddings, whereas the endpoint-only controls retained one endpoint. The endpoint-ablation block used a holdout-weighted ensemble of LogisticRegression, fixed LightGBM, and a PyTorch multilayer perceptron, with weights selected within each outer-training fold. These rows are therefore interpreted as a separate ablation block rather than as a strict nested comparison with the current full-input reference.

For DeepGNHV, mean-pooled ProtT5-derived embeddings represented the human and viral proteins. We evaluated both the full-input representation defined above and a two-endpoint concatenation of the human and viral embeddings. The latter remains a full-input relation baseline because it includes both endpoints; it is not an endpoint-only control. Human-only and virus-only representations were evaluated separately. The verified pipelines used AutoGluon with a gradient-boosting-focused search.

For SAGEPhos, the combined reference used verified ESM2-derived kinase, substrate, and site/window features. Restricted pLM controls retained the site/window, kinase, or full-substrate component. Non-pLM local controls used an 11-residue one-hot window, amino-acid property features, or local k-mer composition. The verified control pipeline used AutoGluon.

Exact pretrained model identifiers, source repositories, and configuration settings are documented in the accompanying repository. Third-party pretrained weights, source benchmark datasets, and other restricted upstream resources are not redistributed; they should be obtained from their original sources under the corresponding licenses. Where redistribution is not permitted, we provide derived split manifests, feature-construction metadata, configuration files, predictions, and result tables sufficient to audit the reported analyses.

### 4.4 Train-derived degree and endpoint-prior controls

The clean MetaESI endpoint-prior feature set used no sequence information. It contained E3 and substrate total counts, positive counts, Laplace-smoothed empirical positive rates,

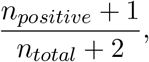

unseen-endpoint indicators, the product and absolute difference of the two smoothed rates, and the log-transformed product of endpoint counts. Two self-label-excluding constructions were evaluated. In the leave-one-edge-out variant, the current training row was removed when computing its endpoint statistics. In the inner-out-of-fold variant, the outer training set was divided into five stratified inner folds with seed 42, and each training row received statistics computed from the other inner folds. Validation and test features were computed from the complete outer training set only. The inner-out-of-fold LogisticRegression result was designated as the primary reported analysis; a fixed HistGradientBoosting classifier and the leave-one-edge-out construction were used as sensitivity analyses. The clean run did not regenerate pLM embeddings, train the original model, use AutoGluon, or tune on the test set.

For DeepGNHV and SAGEPhos we additionally implemented clean cross-fitted train-derived prior diagnostics. DeepGNHV used the original VF1 pair catalog rather than the cache-filtered pLM baseline population, and its features comprised human and virus endpoint counts, positive counts, smoothed positive rates, unseen indicators, and simple interactions. SAGEPhos used kinase, substrate, and exact 11-mer/site-window counts, positive counts, smoothed positive rates, unseen indicators, and simple interactions. Across the eight clean DeepGNHV/SAGEPhos reruns, training-row features excluded the current row’s label by either five-fold inner out-of-fold construction or leave-one-row-out construction; validation and test features used only outer-training labels. These clean prior diagnostics used no exact pair identity or pair frequency and no sequence embeddings.

### 4.5 Restriction-matched MetaESI selector control

For the selector-control block, frozen ESM2 token representations were pooled over either GARD-selected positions or count-matched random positions. Random selections preserved the number of retained tokens for each protein and endpoint role. In the random-both condition, E3 and substrate positions were randomized separately using their respective GARD-selected counts; random-E3 and random-substrate conditions randomized one endpoint while retaining the GARD-pooled representation for the other. All conditions used the same pair-feature construction, split-generation procedure, and fixed LightGBM classifier.

For the row-level selector comparison, random controls were repeated across 10 random seeds and five folds, yielding 50 seed-fold evaluations per random condition; the GARD-selected condition was evaluated across five folds. For E3-cold and substrate-cold selector diagnostics, random controls used five random seeds and five folds, yielding 25 seed-fold evaluations per random condition. This block tests selector attribution within a fixed frozen-pooling pipeline and is not a reimplementation of the complete MetaESI architecture.

### 4.6 Endpoint-cold diagnostics

MetaESI E3-cold and substrate-cold diagnostics were obtained from the verified ESM2 feature-ablation artifacts. DeepGNHV cold-human, cold-virus, and cold-both splits held out the corresponding endpoint identities and used the two-endpoint concatenated ProtT5 representation with the same AutoGluon framework. The corresponding official VF1 two-endpoint result served as the matched warm reference. Cold-split composition and prevalence were reported separately because they differ from the warm split. No SAGEPhos cold split was available in the audited artifacts.

### 4.7 Train-label-shuffle sanity controls

For each task, training labels were randomly permuted with seed 42 while the feature matrices, split assignments, and test labels were retained. A linear SGD classifier with log loss was then fit to the shuffled training labels. The resulting AUROC and AUPRC were used only as basic pipeline sanity checks. They were not interpreted as proof that the original data or evaluation were free from all forms of leakage or dependence.

### 4.8 Metrics and reporting

We report test AUROC and AUPRC. Because AUPRC depends on class prevalence, task-level interpretation was anchored to the corresponding test positive rate, and metrics were not compared across tasks as a ranking. Absolute metrics were prioritized over raw control-to-full ratios. Where source artifacts contained fold- or seed-level variability, the corresponding summary statistics were retained for figures and supplementary tables. No statistical-equivalence conclusion was drawn from similar point estimates or overlapping error bars.

### 4.9 Provenance and reproducibility

Each manuscript result was linked to a repository-verified source file. The clean MetaESI endpoint-prior experiment includes metrics, configuration, feature and split manifests, provenance notes, validation and test predictions, and per-model/per-variant metric files. Equivalent source indices were assembled for the full-input, endpoint, degree-prior, selector-control, cold-split, and label-shuffle blocks. Older MetaESI degree-only results from a superseded protocol were archived and excluded from the manuscript’s primary evidence. Figure source tables were generated directly from the synchronized audited result table.

## Supporting information

Supplementary Data 1. Complete performance-attribution metrics

## 5 Data and Code Availability

Code, configuration files, feature manifests, split manifests, validation/test predictions, synchronized result tables, figure source tables, and the provenance index are available at esm research. Benchmark data remain subject to the access conditions and licenses of the corresponding source projects. Derived split identifiers and processed features are shared where those conditions permit. Exact source identifiers and provenance records for reported results are documented in the accompanying repository.

## 6 Declarations

### 6.1 Author Contributions

Kehan Zhu conceived the study, designed the analysis framework, performed data curation and formal analysis, developed and audited the computational workflows, generated the figures and tables, and wrote the manuscript. Wenbo Zhao contributed exploratory baseline analyses that informed the development of the audit framework. Yuxuan Zhang contributed replication-oriented computational work that informed the project’s methodological development. Zanxian Xia provided scientific oversight, critically reviewed the study framing and manuscript, and approved the final version.

### 6.4 Ethics Statement

This study involved secondary computational analysis of publicly available benchmark resources. No new human participant data, animal experiments, or clinical samples were generated.

