## Supplementary Materials for "Performance Attribution in pLM-Based Biological Relation Prediction"

This supplementary file documents the source-backed methods, sensitivity analyses, diagnostics, figures, and tables supporting the study. It is descriptive by design; the main manuscript provides the primary framing and interpretation.

Unless stated otherwise, AUROC and AUPRC are evaluated on the task-specific held-out test rows. AUPRC should be interpreted relative to the positive prevalence of the evaluated test set. Results from different tasks, row populations, feature spaces, or downstream learners are not treated as directly comparable contribution estimates.

#### 1 Supplementary Methods

##### 1.1 Supplementary Note 1. Construction of self-label-excluding train-derived prior controls

The clean MetaESI endpoint-prior control used no protein sequence embeddings or explicit pair-identity features. Its features comprised E3 and substrate occurrence counts, positive counts, Laplace-smoothed empirical positive rates, unseen-endpoint indicators, and simple cross-endpoint interactions. Laplace-smoothed empirical endpoint positive rates were calculated as

$$\frac{n_{\text{positive}} + 1}{n_{\text{total}} + 2}.$$

Two training-row label-exclusion procedures were evaluated. In the primary inner out-of-fold construction, outer-training rows were divided into five stratified inner folds, and each training row received endpoint statistics estimated from the remaining inner folds. In the leave-one-edge-out construction, the current training row was excluded when computing its endpoint statistics. Validation and test features were derived only from the corresponding outer-training rows.

The primary MetaESI prior-control result used LogisticRegression with balanced class weighting. Fixed HistGradientBoosting and leave-one-edge-out LogisticRegression were evaluated as sensitivity analyses. DeepGNHV and SAGEPhos prior diagnostics were constructed analogously using

cross-fitted training-derived features. DeepGNHV priors used human and virus endpoint statistics, whereas SAGEPhos priors included kinase, substrate, and exact local site-window statistics. Accordingly, the SAGEPhos controls quantify site-window-inclusive prior signal rather than generic degree alone.

##### 1.2 Supplementary Note 2. Restriction-matched MetaESI GARD selector controls

The MetaESI selector-control analysis evaluated a fixed frozen-token pooling pipeline rather than the complete MetaESI architecture. Frozen ESM2 token representations were pooled over either GARD-selected positions or count-matched random positions. Pair features were constructed from the two pooled endpoint representations, their element-wise absolute difference, and their element-wise product. A fixed LightGBM classifier was used for the principal selector-control analyses.

Random controls preserved the number of retained tokens for each protein and endpoint role. In the random-both condition, E3 and substrate positions were randomized separately using their respective GARD-selected counts. In the random-E3 and random-substrate conditions, one endpoint was randomized while the GARD-pooled representation was retained for the other endpoint.

For row-level evaluation, random controls were repeated across 10 random seeds and five folds, yielding 50 seed-fold evaluations per random condition; the GARD-selected condition was evaluated across five folds. For E3-cold and substrate-cold selector diagnostics, random controls were repeated across five random seeds and five folds, yielding 25 seed-fold evaluations per random condition. Reported variability values are source-summary standard deviations. These analyses assess whether selected token identity provides an observed predictive advantage beyond a matched restriction of the frozen representation; they do not establish formal equivalence, evaluate the full MetaESI architecture, or determine the biological relevance of individual GARD-selected regions.

##### 1.3 Supplementary Note 3. Endpoint-cold and train-label-shuffle diagnostics

Endpoint-cold analyses were used as supplementary stress tests for endpoint novelty. In MetaESI, E3-cold and substrate-cold diagnostics were obtained from a saved ESM2 feature-ablation block and are interpreted separately from the current full-input rerun. In DeepGNHV, cold-human, cold-virus, and cold-both evaluations used the two-endpoint frozen representation under matched warm and cold evaluation settings. No kinase-cold, substrate-cold, or site-cold diagnostic was available for SAGEPhos.

These endpoint-cold analyses do not constitute hard-negative, matched-negative, family-aware, homology-aware, or external-cohort evaluation. They assess performance under the available endpoint-novelty splits only.

For train-label-shuffle sanity controls, training labels were randomly permuted while feature matrices, split assignments, and test labels were retained. The resulting AUROC and AUPRC values were used solely to confirm that the evaluated control pipelines lost useful training-label association after shuffling. These diagnostics do not establish the absence of all possible leakage or benchmark-dependence mechanisms.

Detailed source data, configuration files, split manifests, feature manifests, prediction files, and provenance records are available in the accompanying repository.

### 2 Supplementary Figures

#### 2.1 Figure S1. Endpoint-cold diagnostics

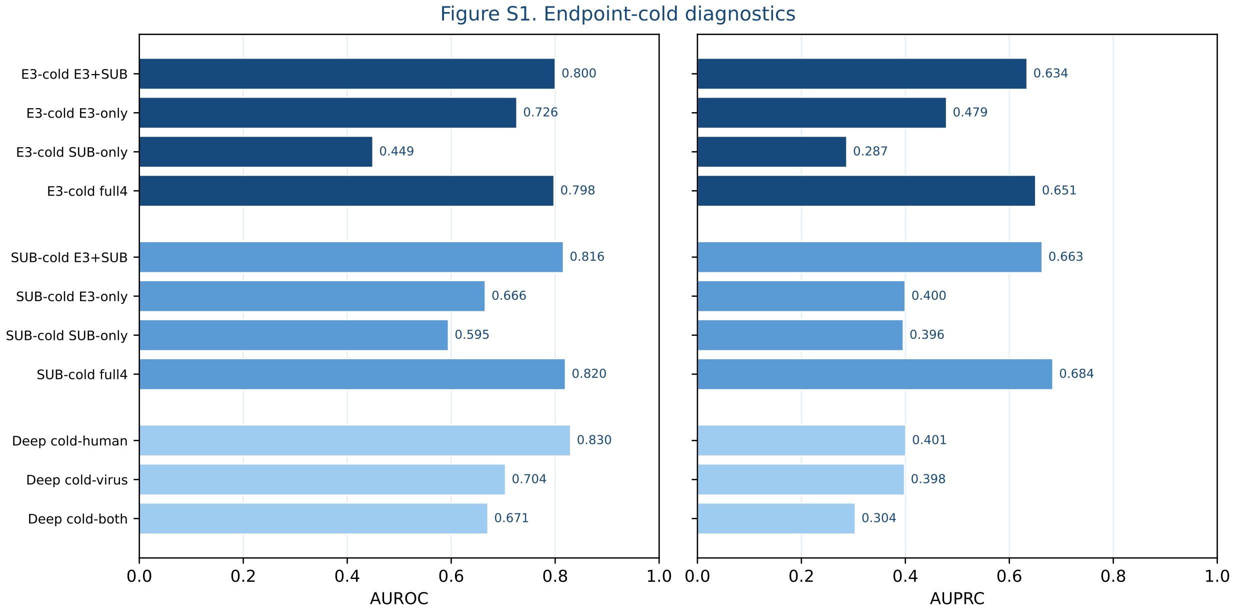


Figure S1: Endpoint-cold diagnostics for MetaESI and DeepGNHV. Bars show AUROC and AUPRC under the available endpoint-cold evaluation settings. MetaESI includes E3-cold and substrate-cold diagnostics for frozen full-input, two-endpoint, E3-only, and substrate-only ESM2 feature settings derived from a separate feature-ablation block. DeepGNHV includes cold-human, cold-virus, and cold-both evaluation using the two-endpoint frozen representation. These analyses assess endpoint novelty under the available split definitions and are not hard-negative, matched-negative, family-aware, or homology-aware evaluations.

#### 2.2 Figure S2. Train-label-shuffle sanity controls

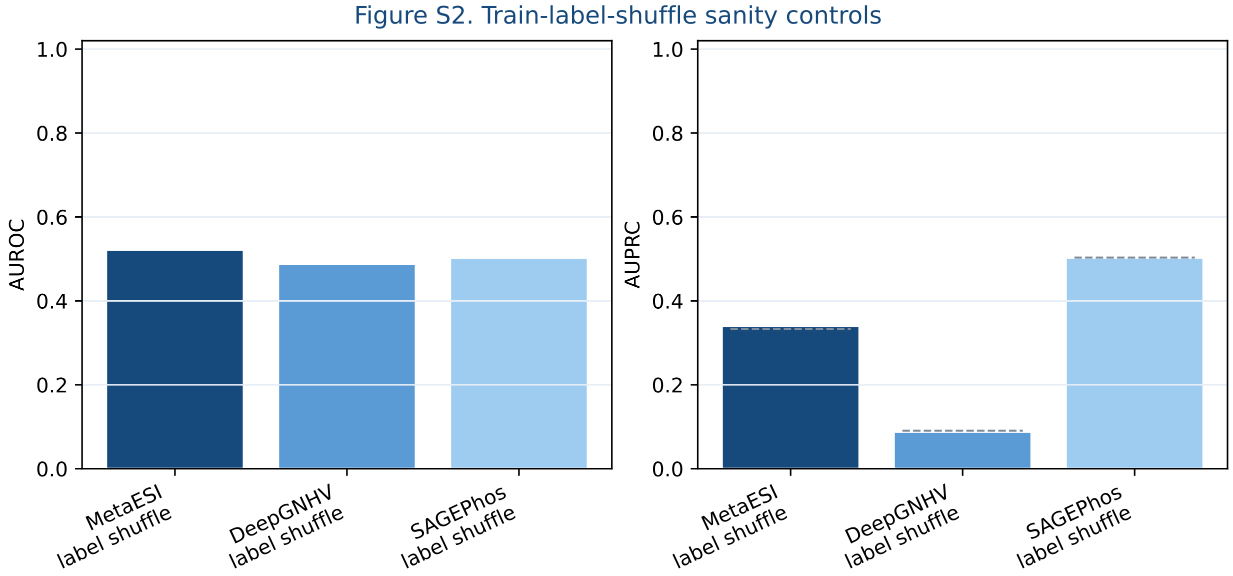


Figure S2: Train-label-shuffle sanity controls across MetaESI, DeepGNHV, and SAGEPhos. Training labels were permuted while feature matrices, split assignments, and test labels were retained. AUROC values returned to approximately 0.5, and AUPRC values tracked the corresponding positive prevalence of each evaluated test set, indicated by dashed lines. These results are pipeline sanity checks and do not establish the absence of all possible forms of leakage or benchmark dependence.

#### 2.3 Figure S3. MetaESI GARD selector and coverage diagnostics

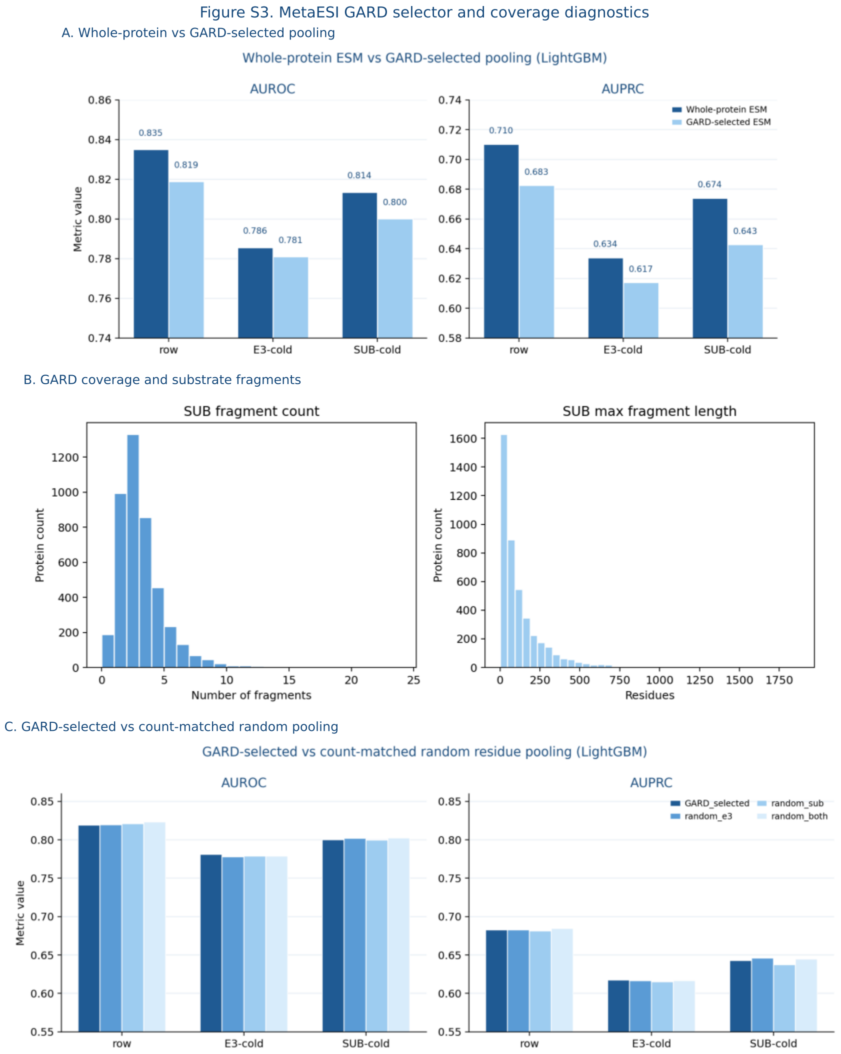


Figure S3: MetaESI GARD selector and coverage diagnostics. Panel A compares whole-protein frozen ESM pooling with GARD-selected frozen pooling across row-level, E3-cold, and substrate-cold settings. Panel B summarizes substrate-side GARD fragmentation through the number of retained fragments per protein and the maximum retained fragment length. Panel C compares GARD-selected pooling with count-matched random pooling across row-level, E3-cold, and substrate-cold settings. Random row-level controls were evaluated across 10 random seeds and five folds; endpoint-cold random controls were evaluated across five random seeds and five folds; GARD-selected pooling was evaluated across five folds. These panels provide selector-control and coverage context for the fixed frozen-pooling analysis and do not evaluate the complete MetaESI architecture.

### 3 Supplementary Tables

#### 3.1 Table S1. MetaESI clean endpoint-prior sensitivity variants

Table S1: MetaESI clean endpoint-prior sensitivity variants.

| Feature construction | Classifier | AUROC | AUPRC | Accuracy | Balanced accuracy | F1 | MCC |
|---|---|---|---|---|---|---|---|
| leave one edge out | LogisticRegression | 0.849 | 0.692 | 0.766 | 0.791 | 0.712 | 0.550 |
| leave one edge out | HistGradientBoosting fixed | 0.699 | 0.572 | 0.708 | 0.642 | 0.504 | 0.307 |
| inner oof | LogisticRegression | 0.846 | 0.676 | 0.746 | 0.779 | 0.698 | 0.527 |
| inner oof | HistGradientBoosting fixed | 0.847 | 0.684 | 0.759 | 0.776 | 0.696 | 0.522 |

MetaESI self-label-excluding endpoint-prior sensitivity analyses. The table compares inner out-of-fold and leave-one-edge-out feature construction using LogisticRegression and fixed HistGradientBoosting. Features comprise train-derived endpoint counts, positive counts, Laplace-smoothed empirical positive rates, unseen-endpoint indicators, and simple cross-endpoint interactions; no sequence embeddings are used. The primary manuscript result is the inner out-of-fold LogisticRegression model. Remaining rows are sensitivity analyses.

#### 3.2 Table S2. Endpoint-cold diagnostic metrics and split composition

Table S2: Endpoint-cold diagnostic metrics and split composition.

| Task | Feature set | Split | Classifier | AUROC | AUPRC | Test n | Test prevalence |
|---|---|---|---|---|---|---|---|
| MetaESI | E3+SUB | E3-cold | Holdout-weighted ens. | 0.800 | 0.634 | – | – |
| MetaESI | E3-only | E3-cold | Holdout-weighted ens. | 0.726 | 0.479 | – | – |
| MetaESI | SUB-only | E3-cold | Holdout-weighted ens. | 0.449 | 0.287 | – | – |
| MetaESI | mvp_full4 | E3-cold | Holdout-weighted ens. | 0.798 | 0.651 | – | – |
| MetaESI | E3+SUB | SUB-cold | Holdout-weighted ens. | 0.816 | 0.663 | – | – |
| MetaESI | E3-only | SUB-cold | Holdout-weighted ens. | 0.666 | 0.400 | – | – |
| MetaESI | SUB-only | SUB-cold | Holdout-weighted ens. | 0.595 | 0.396 | – | – |
| MetaESI | mvp_full4 | SUB-cold | Holdout-weighted ens. | 0.820 | 0.684 | – | – |
| DeepGNHV | concat_hv | cold_human | WE L2 | 0.830 | 0.401 | 44716.000 | 0.088 |
| DeepGNHV | concat_hv | cold_virus | WE L2 | 0.704 | 0.398 | 48437.000 | 0.115 |
| DeepGNHV | concat_hv | cold_both | WE L2 | 0.671 | 0.304 | 15660.000 | 0.111 |

Endpoint-cold diagnostic metrics for MetaESI and DeepGNHV. MetaESI rows report E3-cold and substrate-cold diagnostics from a separate ESM2 feature-ablation block. DeepGNHV rows report cold-human, cold-virus, and cold-both evaluation using the two-endpoint frozen representation, together with test-set size and positive prevalence where available. These analyses assess endpoint novelty under the available split definitions and are not hard-negative, matched-negative, family-aware, or homology-aware evaluations.

##### 3.3 Table S3. MetaESI GARD-selected and count-matched random pooling summary

Table S3: MetaESI GARD-selected and count-matched random pooling summary.

| Split | Model | Selector mode | AUROC mean (SD) | AUPRC mean (SD) | Seeds | Runs/folds | MCC |
|---|---|---|---|---|---|---|---|
| E3-cold | LightGBM | GARD selected | 0.781 (0.026) | 0.617 (0.038) | 1 | 5 | 0.259 |
| E3-cold | LightGBM | random both | 0.779 (0.024) | 0.617 (0.045) | 5 | 25 | 0.277 |
| E3-cold | LightGBM | random e3 | 0.778 (0.024) | 0.617 (0.034) | 5 | 25 | 0.277 |
| E3-cold | LightGBM | random sub | 0.779 (0.023) | 0.615 (0.039) | 5 | 25 | 0.273 |
| E3-cold | LR | GARD selected | 0.721 (0.023) | 0.502 (0.028) | 1 | 5 | 0.291 |
| E3-cold | LR | random both | 0.732 (0.022) | 0.519 (0.031) | 5 | 25 | 0.312 |
| E3-cold | LR | random e3 | 0.715 (0.023) | 0.501 (0.029) | 5 | 25 | 0.290 |
| E3-cold | LR | random sub | 0.729 (0.019) | 0.515 (0.027) | 5 | 25 | 0.305 |
| SUB-cold | LightGBM | GARD selected | 0.800 (0.010) | 0.643 (0.017) | 1 | 5 | 0.352 |
| SUB-cold | LightGBM | random both | 0.802 (0.014) | 0.645 (0.021) | 5 | 25 | 0.361 |
| SUB-cold | LightGBM | random e3 | 0.802 (0.010) | 0.646 (0.019) | 5 | 25 | 0.365 |
| SUB-cold | LightGBM | random sub | 0.800 (0.014) | 0.638 (0.020) | 5 | 25 | 0.355 |
| SUB-cold | LR | GARD selected | 0.735 (0.014) | 0.508 (0.012) | 1 | 5 | 0.303 |
| SUB-cold | LR | random both | 0.732 (0.013) | 0.515 (0.018) | 5 | 25 | 0.300 |
| SUB-cold | LR | random e3 | 0.735 (0.009) | 0.513 (0.015) | 5 | 25 | 0.310 |
| SUB-cold | LR | random sub | 0.732 (0.011) | 0.514 (0.019) | 5 | 25 | 0.291 |
| row | LightGBM | GARD selected | 0.819 (0.007) | 0.683 (0.011) | 1 | 5 | 0.416 |
| row | LightGBM | random both | 0.823 (0.010) | 0.684 (0.015) | 10 | 50 | 0.425 |
| row | LightGBM | random e3 | 0.820 (0.007) | 0.683 (0.011) | 10 | 50 | 0.422 |
| row | LightGBM | random sub | 0.821 (0.009) | 0.682 (0.015) | 10 | 50 | 0.418 |
| row | LR | GARD selected | 0.753 (0.018) | 0.533 (0.027) | 1 | 5 | 0.350 |
| row | LR | random both | 0.756 (0.013) | 0.539 (0.018) | 10 | 50 | 0.358 |
| row | LR | random e3 | 0.754 (0.015) | 0.535 (0.025) | 10 | 50 | 0.353 |
| row | LR | random sub | 0.758 (0.010) | 0.542 (0.016) | 10 | 50 | 0.358 |

MetaESI GARD-selected and count-matched random pooling results across row-level, E3-cold, and substrate-cold settings. The table reports mean AUROC, AUPRC, MCC, source-summary standard deviations, selector mode, classifier, random-seed count, and number of fold-level evaluations. GARD-selected pooling uses one fixed selected-region definition evaluated across five folds. Row-level random controls use 10 random seeds across five folds, whereas endpoint-cold random controls use five random seeds across five folds. This table summarizes a fixed frozen-token selector-control pipeline and does not evaluate the complete MetaESI architecture.

#### 4 Supplementary Data

##### 4.1 Supplementary Data 1. Complete performance-attribution metrics

A machine-readable CSV file is provided separately.
